# Comparative sub-cellular proteome analyses reveals metabolic differentiation and production of effector-like molecules in the dieback phytopathogen *Phytophthora cinnamomi*

**DOI:** 10.1101/2022.02.15.480627

**Authors:** Christina E. Andronis, Silke Jacques, Richard Lipscombe, Kar-Chun Tan

**Affiliations:** Centre for Crop and Disease Management, Curtin University, Bentley, WA, Australia; Proteomics International, Nedlands, WA, Australia

**Keywords:** Oomycete, *Phytophthora cinnamomi*, dieback, proteome, zoospore, secretome, virulence, effectors

## Abstract

Phytopathogenic oomycetes pose a significant threat to global biodiversity and food security. The proteomes of these oomycetes likely contain important factors that contribute to their pathogenic success, making their discovery crucial for elucidating pathogenicity. Oomycetes secrete effector proteins that overcome or elicit a defence response in susceptible hosts. *Phytophthora cinnamomi* is a root pathogen that causes dieback in a wide variety of crops and a range of native vegetation world-wide. Virulence proteins produced by *P. cinnamomi* are not well defined and a large-scale approach to understand the biochemistry of this pathogen has not been documented. Here, soluble mycelial, zoospore and secreted proteomes were obtained and label-free quantitative proteomics was used to compare the composition of protein content of the three sub-proteomes by matching the MS/MS data to a sequenced *P. cinnamomi* genome. Mass spectra matched to a total of 4635 proteins, validating 17.7% of the predicted gene set of the *P. cinnamomi* genome. The mycelia were abundant in transporters for nutrient acquisition, metabolism and cellular proliferation. The zoospores had less metabolic related ontologies than the mycelia but were abundant in energy generating, motility and signalling associated proteins. Virulence-associated proteins were identified in the secretome such as candidate effector and effector-like proteins, which interfere with the host immune system. These include hydrolases, cell wall degrading enzymes, putative necrosis-inducing proteins and elicitins. The secretome elicited a hypersensitive response on the roots of a model host and thus suggests evidence of effector activity.

**Significance:** *Phytophthora cinnamomi* is a phytopathogenic oomycete that causes dieback disease in native vegetation and several horticultural crops such as avocado, pineapple and macadamia. Whilst this pathogen has significance world-wide, its pathogenicity and virulence have not been described in depth. We carried out comparative label-free proteomics of the mycelia, zoospores and secretome of *P. cinnamomi*. This study highlights the differential metabolism and cellular processes between the sub-proteomes. Proteins associated with metabolism, nutrient transport and cellular proliferation were over represented in the mycelia. The zoospores have a specialised proteome showing increased energy generation geared towards motility. Candidate effectors and effector-like secreted proteins were also identified, which can be exploited for genetic resistance. This demonstrates a better understanding of the biology and pathogenicity of *P. cinnamomi* infection that can subsequently be used to develop effective methods of disease management

## Introduction

*Phytophthora cinnamomi* is a soil-borne phytopathogenic oomycete that causes significant economic and environmental losses world-wide. In Australia, the pathogen causes dieback on native vegetation species as well as major horticultural crops such as avocados, macadamia, pineapple and stone fruits ^1^. There is no evidence of genetic resistance being used to combat dieback disease, hence it is largely controlled by the use of the oomyceticide phosphite which can only slow down the progression of disease severity and spread ^2^. The mode of action of phosphite is not well understood and the application of this chemical to crops and vegetation is highly labour intensive as it is injected manually into the trunks of susceptible trees. There is emerging resistance to phosphite particularly in avocado orchards ^3^. Therefore, there is a need to better understand the pathogenicity of this system to develop more sustainable methods of combatting dieback disease.

The lifecycle of *P. cinnamomi* is characterised by several forms including vegetative and reproductive stages ^4^. Briefly, vegetative mycelia grow within host tissue where they mature into sexual fruiting bodies called sporangia which in turn produce motile sexual cells called zoospores. Zoospores travel through watery soils with the aid of flagellum until they encounter the roots of a susceptible host ^5^. Once contact is achieved, the zoospores begin to secrete an arsenal of proteins to initiate the formation dedicated structures. A protective cyst forms on the surface of the root tip from which sprout appressoria that invade the plants tissue, transporting effectors and nutrients between host and pathogen.

Transcriptomics is the method of choice for comparative gene expression analysis in many studies ^6^. In contrast to the transcriptome, the proteome can be used to identify what is present and abundant in different life stages, tissues and cellular compartments of the pathogen ^7^. Proteomics has been used to capture such virulence factors in other oomycetes ^8,9^. Virulence factors such as effectors are proteins that are released by the pathogen and assist in infection by damaging host tissue or interfering with host immunity. In oomycetes, effectors include necrosis inducing proteins, RXLR and CRINKLER effectors and elicitins ^10^.

The protein content of *P. cinnamomi* has not been previously studied and virulence factors that contribute to its pathogenic success remain unknown. The present study used mass spectrometry-based proteomics to elucidate the biochemistry of three sub-proteomes of *P. cinnamomi.* The mycelia, secretome and zoospore sub-proteomes were qualitatively and quantitatively analysed to gain an understanding of which key factors are abundant that contribute to the virulence of this pathogen. Furthermore, a cell-free secretome preparation demonstrated necrosis on lupin (*Lupinus angustifolia*) roots suggesting the presence of necrosis inducing factors. The secretome was further profiled to determine the composition of effector-like proteins that may function to dictate the outcome of infection. By studying each of these stages of growth separately and identifying these virulence-associated proteins, we provide biochemical snapshot of the organism and identify important factors that contribute to pathogenicity.

## Experimental procedures

### *Phytophthora* culture and preparation

*Phytophthora cinnamomi* (MU94-48) cultures were obtained from the Centre of Phytophthora Science and Management (Murdoch University, Australia) and maintained by passaging through *Malus domestica* cv. Granny Smith fruit ^11,12^. Mycelia were grown on V8 juice (Campbells) agar with growth periods of 5-7 days in the dark at room temperature. Mycelia were scraped off the surface of the plates and placed into Erlenmeyer flasks containing 40 mL of Ribeiros Minimal Media ^13^. Liquid cultures were incubated for 3 days in the dark at room temperature. Mycelia were harvested by centrifugation at 4000g for 30 minutes and washed twice with water. The liquid media was passed through a 0.22 µm filter to remove mycelial fragments and obtain the secretome. Zoospores were produced as previously described ^12^. Purified mycelia, secretome and zoospores were snap-frozen in liquid nitrogen and freeze-dried. All sub-proteomes were prepared in triplicate.

### Protein extraction and digestion

Dried mycelia and zoospores were ground to a fine powder using beads and Biosprint shaker (Qiagen). 300 µl of extraction buffer (25 mM Tris-HCl pH 7.5, 0.25% SDS, 50 mM Na2PO4, 1 mM Na2F, 50 µM Na3VO4 and 1 mM PMSF in the presence of a protease inhibitor cocktail (Sigma)) was added to the total cell lysates and incubated for 30 min on ice with occasional vortexing. Samples were centrifuged at 15000 g for 30 minutes at 4 °C and the supernatant was transferred to a new tube. The dried secretome was resuspended in 3 mL of water and proteins from the three sub-proteomes were precipitated with 6 volumes of acetone. Samples were initially digested with trypsin for 3 hours at 37 °C to assist in solubilising the pellet, reduced and alkylated with 50 mM tris (2-carboxyethyl)phosphine (Thermo Scientific, Waltham) and 200 mM methyl methanethiosulfonate (Sigma, St Louis) respectively. Samples were digested again overnight at 37°C with trypsin (Sigma, St Louis) at a ratio of 1:10, subsequently desalted on a Strata-X 33 um polymeric reverse phase column (Phenomenex, Torrance, CA, USA) and dried in a vacuum centrifuge.

### Mass spectrometry

Samples were analysed by electrospray ionisation mass spectrometry using a Thermo UltiMate 3000 nanoflow UHPLC system (Thermo Scientific) coupled to a Q Exactive HF mass spectrometer (Thermo Scientific). Approximately 1 µg of peptides were loaded onto an Acclaim™ PepMap™ 100 C18 LC Column, 2 µm particle size x 150 mm (Thermo Scientific) and separated with a linear gradient over 190 minutes of water/acetonitrile/0.1% formic acid (v/v).

### Qualitative, quantitative and functional analysis

Label-free quantification was performed using the Proteome Discoverer 2.3 using the label-free precursor quantification workflow template ^14^. Spectra were matched against the *P. cinnamomi* MU94-48 genome ^12^. Sub-proteomes were relatively quantified using the default precursor ion quantifier. For Identification, proteins with one peptide identified were used and for quantification proteins with two or more peptides were used. The FDR was set at <1. The criteria used for significant differential abundance were p < 0.05. Gene ontologies were assigned using Interpro Scan 86.0. Gene Ontology enrichment within and between sub-proteomes was determined using Fisher’s exact test. To further examine protein function, KEGG orthologues and Interpro annotations were used.

### Necrosis induction assay

The cell-free secretome used for inoculation of lupin seedlings was prepared as above using acetone precipitation to remove non-proteinaceous molecules. The protein pellet of the secretome was resuspended in 3 mL of water and left to solubilise for one hour. The protein content was measured using a nanodrop and concentration was made up to 8 mg/mL and 500 µl of the secretome was used. A control was prepared with the growth media and an additional control containing 8 mg/mL of BSA was used to ensure the concentrated protein did not cause an effect on the plant. *Lupinus angustifolia* (Tanjil) seeds were germinated by soaking in water and left for three days in a plastic container lined with moist Whatmann paper at room temperature. The seedlings were inoculated with the secretome by immersing the root tips into 1.5 mL Eppendorf tubes containing the secretome. Response from the secretome was measured by scoring the severity of the lesion colour and size, three days after inoculation. EffectorP 3.0, PFAM 34.0 and PHI-Base 4.12 were used on the identified protein sequences to further characterise the composition of the secretome and identify the candidate effectors.

## Results

### Protein identifications in the sub-proteomes

Mycelia, zoospore and secreted proteins were selected for proteomic analysis as they reflect vegetative and motile stages of the pathogen. Using high confidence peptides, 4635 proteins from the *P. cinnamomi* annotation were identified (Table 1). The total protein identifications for each sub-proteomes are shown in Supplementary material 1 and 2. This accounts for 17.7% of the predicted annotated genes in the *P. cinnamomi* genome ^12^. Of these, 1070, 698 and 278 were unique to the mycelia, zoospores and secretome respectively (Figure 1). A high proportion of 1803 accounting for 38.9% of identified proteins was common between the mycelia and zoospores, while 2.2% and 1.2% were common between the mycelia and secretome, and secretome and zoospore respectively. 629 proteins were shared between all three sub-proteomes.

**Table 1.**
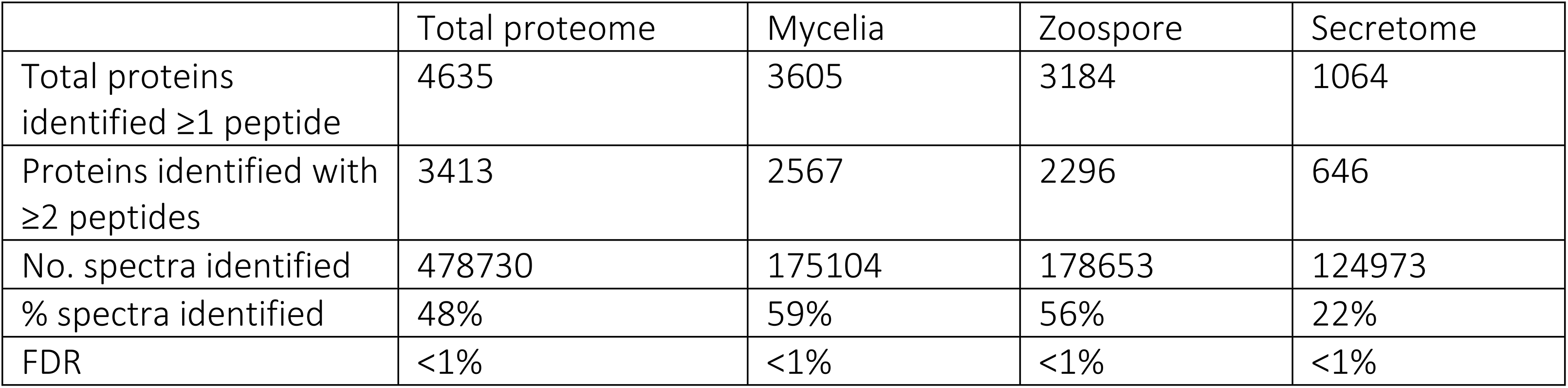
Protein identifications and spectral matches of sub-proteomes

**Figure 1.**
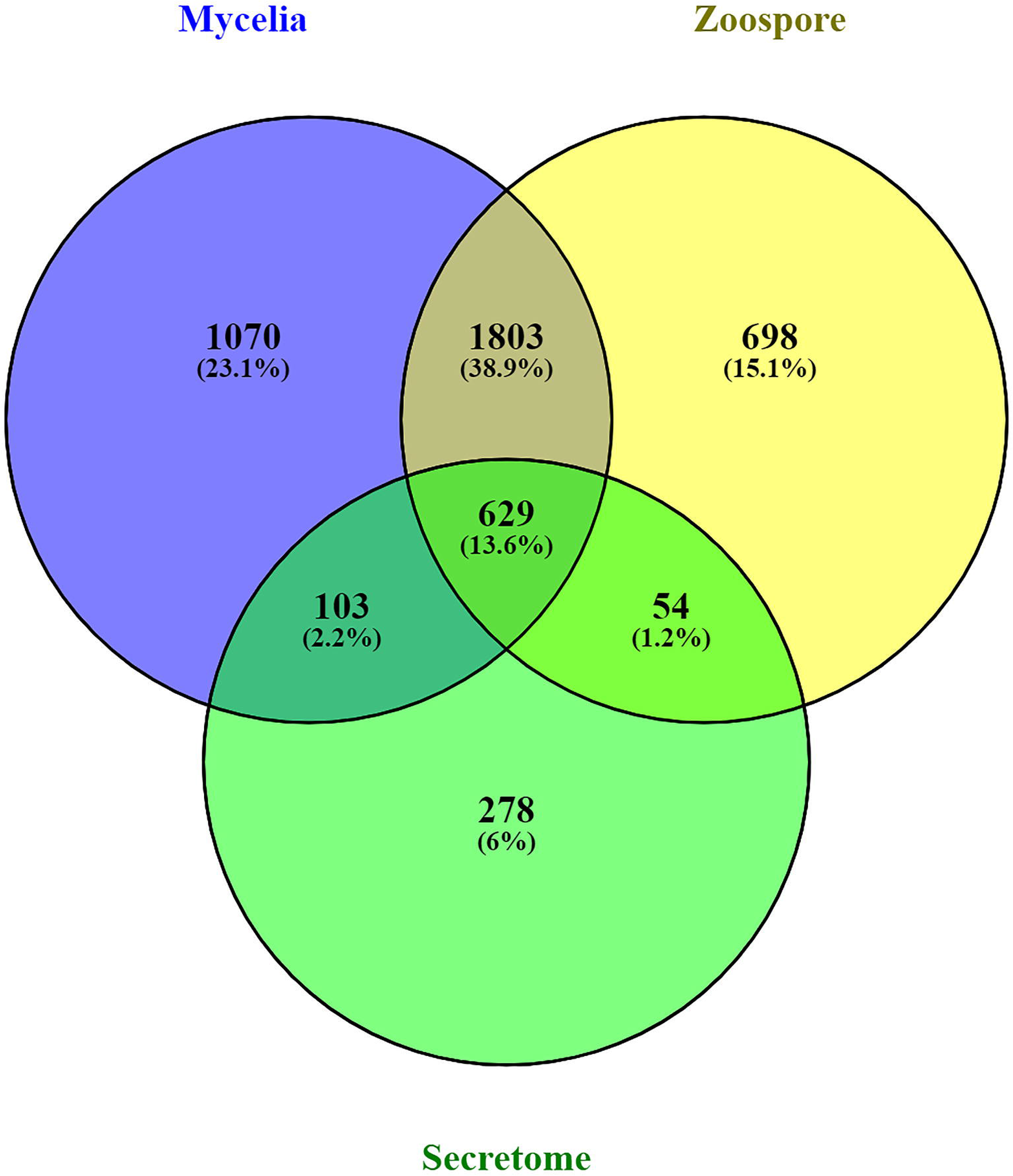
Sub-proteome identification of *Phytophthora cinnamomi* indicating number of unique and common proteins across the mycelia, zoospores and secretome.

### The mycelia metabolism focuses on saprotrophy

The mycelia must continually obtain nutrients for saprophytic growth and successful infection of neighbouring host tissue. The functional composition of the 1070 proteins unique to the mycelia was significantly enriched with Gene Ontologies (GO) associated with metabolic processes and cellular functioning for nutrient metabolism and growth (Table 2).Transporters such as amino acid/ polyamine (IPR00229), metal ion transporters (K14686, IPR00852), inositol and sugar transporters (IPR00366), ABC transporters (K05643) and sulphate transmembrane transporters and permeases (GO:0015116, K03321) were found in the mycelia. Sugar, carbohydrate, lipid and amino acid metabolic processes were enriched in the mycelia (GO:0006000, GO:0005975, GO:0006629, GO:0044255, GO:0006520), including phospholipid, glycerol and fatty acid metabolism and biosynthesis (K13356, IPR0212, IPR01312, IPR02228), indicating metabolism of nutrients. Additionally, biosynthesis related to cell proliferation was also abundant in the mycelia (GO:0009058, GO:0007093). These include aminotransferases (K00814), proteins associated with chromosome separation such as condensing (GO:0051304) and DNA synthesis and cell division enzyme including thymidine kinase (K08866). The abundance of these genes suggests that the primary biochemical functions of the mycelia focuses on nutrient acquisition and metabolism during vegetative growth.

**Table 2.**
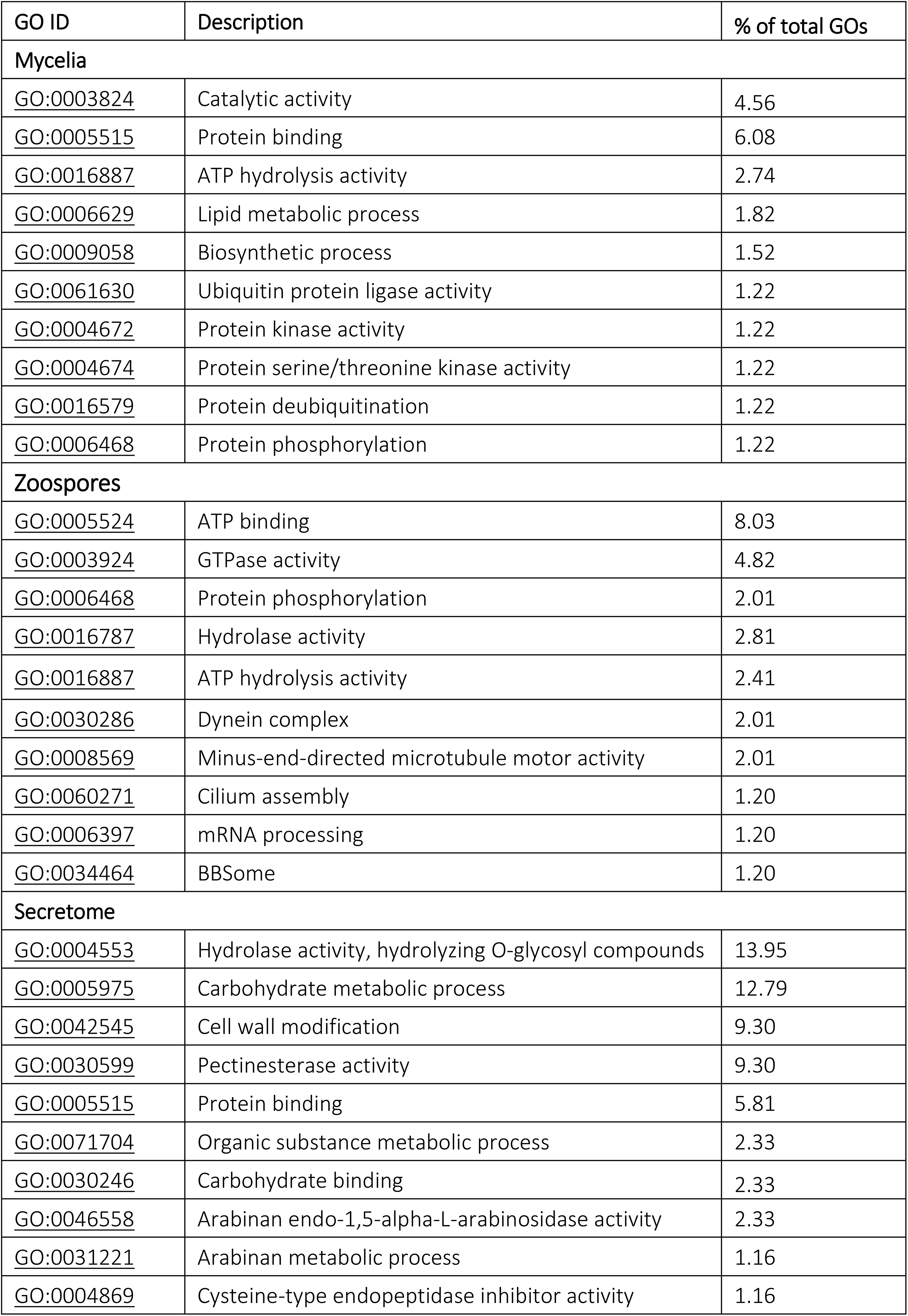
Top 10 significantly enriched GOs of the mycelia, zoospores and extracellular sub-proteomes by fisher’s exact test (p <0.05).

### The zoospore proteome is enriched in proteins that are associated with energy generation, motility and signalling

The zoospores play an important role in the infection process as they enable spread of disease and can initiate the infection process by encysting on plant root tissue. The enriched GOs in the proteins unique to the zoospores were mostly associated with energy production and motility (Table 2). The full list of enriched gene ontologies for these sub-proteomes is shown in Supplementary material 1. Major energy generation and secondary signalling messengers included ATP (GO:0005524, GO:0016887) and GTPases (GO:0003924). Complexes associated with motility include dynein (GO:0030286), cilium assembly (GO:0060271, GO:0030992, GO:0042073), and the BBSome (GO:0034464). The genes associated with these ontologies are domains of the motor complex (K10408, K16746, IPR01104, IPR02878, IPR03272) and constituents of intraflagellar transport and signalling (IPR029600). In addition, putative hydrolases (GO:0016787) were also enriched in the zoospores including glycoside hydrolases (IPR00154) and acid phosphatases (KO22390). Quantitative comparison between the zoospores and mycelia indicated an abundance in proteins associated with RNA/ protein binding and modification (Figure 2). An increase in response to nitrosative stress (GO:0051409) and nitric oxide dioxygenase activity (GO:0008941) suggests preparation to defend against host immunity ^15,16^. These proteins along with the increase in protein phosphorylation play roles in signalling in response to external stimuli.

**Figure 2.**
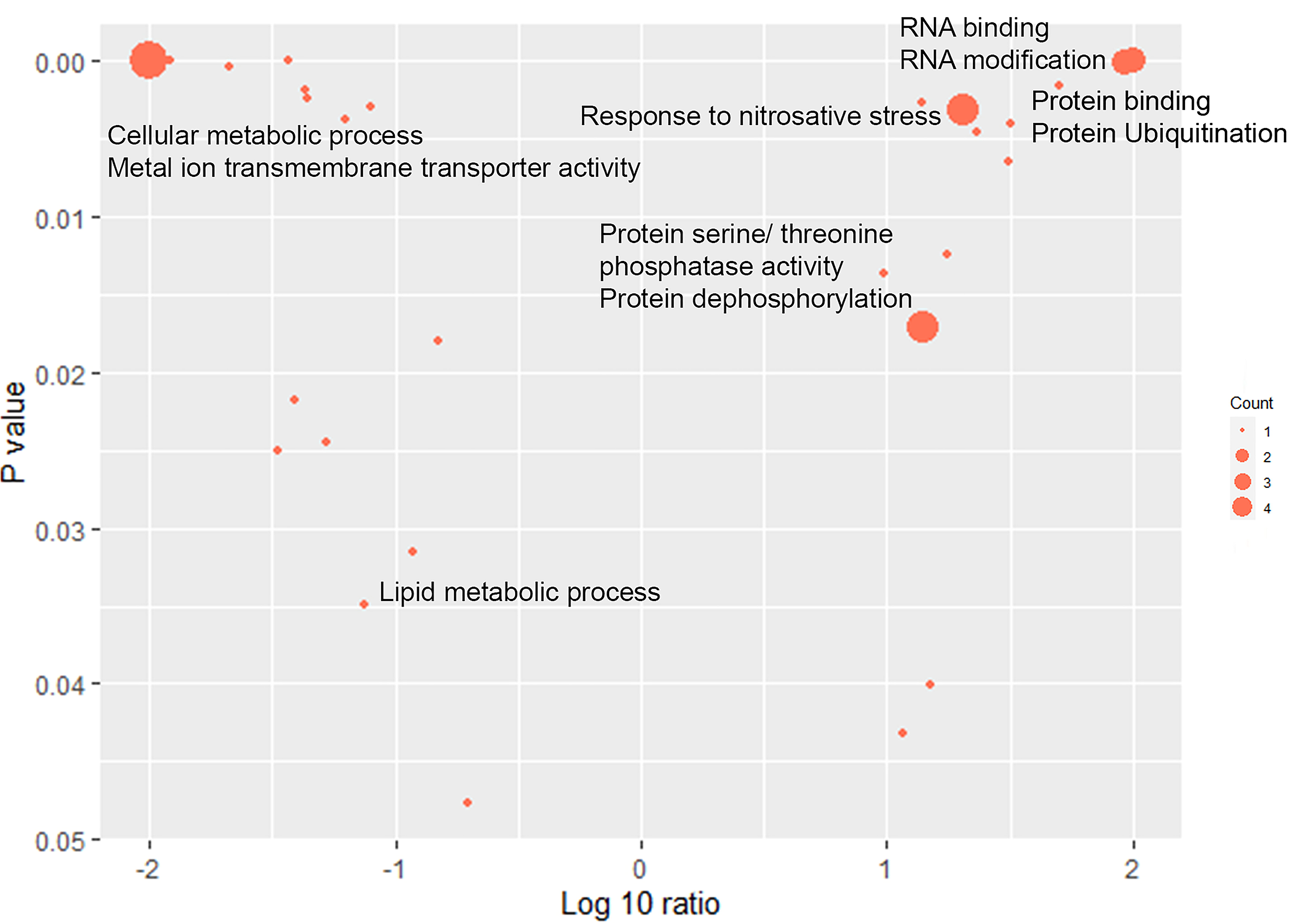
GOs that are over- and under-represented in the zoospore proteome. The X-axis indicates the log 10 ratio of the abundance ratio of zoospores to the mycelia labelled with their associated GOs. The size correlates with the number of genes annotated with each GO. A full list of significantly differentially abundant proteins and their associated GOs are shown in Supplementary material 2.

Quantitative analyses between the zoospores and mycelia also demonstrated that metabolic processes such as cellular and lipid metabolism were down-regulated in the zoospores. Metabolic processes such as lipid metabolism (GO:0006629), amino acid metabolism (GO: GO:0009072) and cellular metabolic processes (GO:0044237) were lower in abundance. RNA binding and modification (GO:0003723, GO:0009451) was over-represented in the zoospores where these genes were involved in post-transcriptional modification, translin transport and ribonucleases (IPR002848, IPR016068). Protein ubiquitination (GO:0016567), phosphorylation (GO:0006470) and protein binding (GO:0005515) were also higher in abundance in the zoospores with genes specific to ubiquitin ligase components such as HECT domains (IPR035983) and SPRY domains (IPR003877) which have important roles in protein activation and signalling. This indicates that zoospores are geared towards energy production to fuel motor activity and assembly of structural components along with rapid response to external stimuli.

### Analysis of the secretome proteome identifies putative effectors and enrichment in CWDEs

In other *Phytophthora* species, the secretome contains a plethora of virulence-associated molecules that interfere with host immunity and assist in successful infection. The mycelia produce this set of proteins to continually degrade host tissue, dampen host immunity and facilitate the spread of disease to neighbouring roots. The secretome was enriched with GOs involved in cell wall degradation and necrosis (Table 2). Cell wall modification proteins associated with structural components of plants such as pectinases (GO:0030599, K01051), arabinose (GO:0046558, GO:0019566, GO:0031221), carbohydrates (GO:0005975) and hydrolases (GO:0016787, GO:0004553) facilitate host penetration by breaking down plant tissue. Pectin associated genes included those involved in pectin lyase folding (IPR012334), catalytic activity (IPR000070), genes acting on active sites (IPR033131) and pectin associated virulence factors (IPR011050). Hydrolases acting on glycosidic bonds (K01184, IPR000743, IPR000490, IPR017853), concanavalin lectins (K20844, IPR013320) and acid phosphatase hydrolases (K22390, IPR008963, IPR041792) were enriched in the secretome. Cysteine-type endopeptidase inhibitors (GO:0004869, IPR000010) were identified in the secretome, which is characteristic of many secreted effector proteins.

To determine if the secretome contained candidate effectors that induce a hypersensitive response in plants, Lupin seedlings were inoculated with a cell free secretome preparation. *P. cinnamomi* MU94-48 infects and causes dark necrotic lesions on *Lupinus angustifolius* roots ^17^. Exposure of the seedlings to the cell free secretome caused a hypersensitive response where dark lesions were developed three days post-inoculation (Figure 3). The secretome was also treated with a protease to confirm that the HR was caused by proteins only, however the protease control caused a lesion response in the root. To identify effectors, the secretome proteome was subjected to the effector prediction software, EffectorP (Table 3). This showed a predicted 422 of the 1064 proteins as cytoplasmic and/or apoplastic effectors. The assigned PFAM domains of the proteins within the secretome demonstrated the presence of several types of effectors including 16 cysteine-rich secreted proteins, 8 necrosis inducing proteins, 14 elicitins and one protein with an RXLR motif (Table 3). When searched against the PHI-Base database, several homologs to effectors in other *Phytophthora* species were found, including elicitins INF1 and INF2A, protease inhibitors such as EPI1 and EPI2, suppressors of elicitor mediated plant defence response such as GIP and several adhesive and necrosis inducing proteins such as PsoNIP, CBEL-GP34, Pc129485 and PcPL. This suggests that the secretome is enriched with molecules with putative functions in host penetration and immune system manipulation.

**Figure 3.**
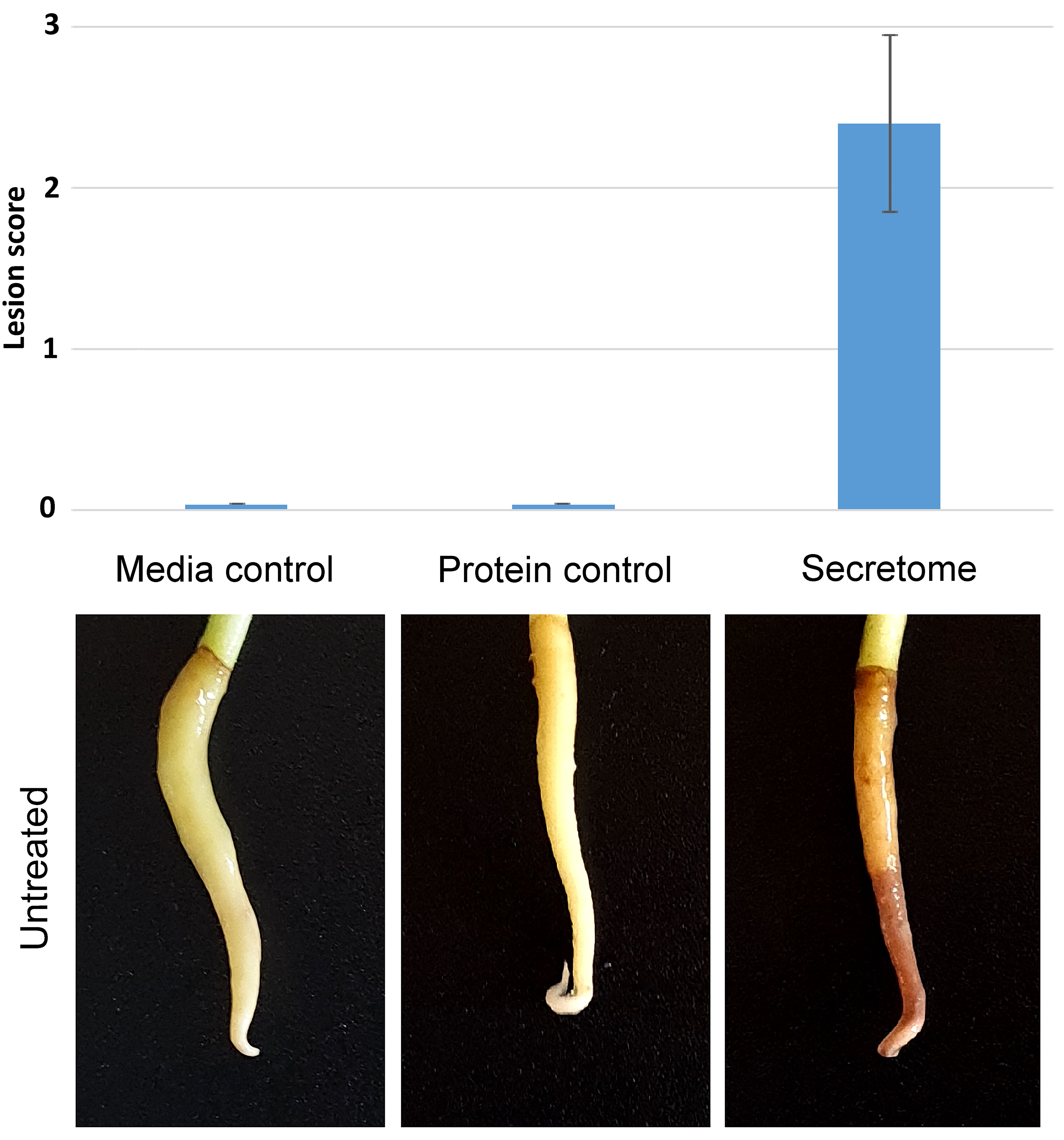
Lesion score and necrosis on *L. angustifolia* roots caused by the cell-free secretome extract of *P. cinnamomi.* A media and a equivalent concentration protein control were used. Lesions were scored from zero to three by the colour and size of necrotic tissue.

**Table 3.**
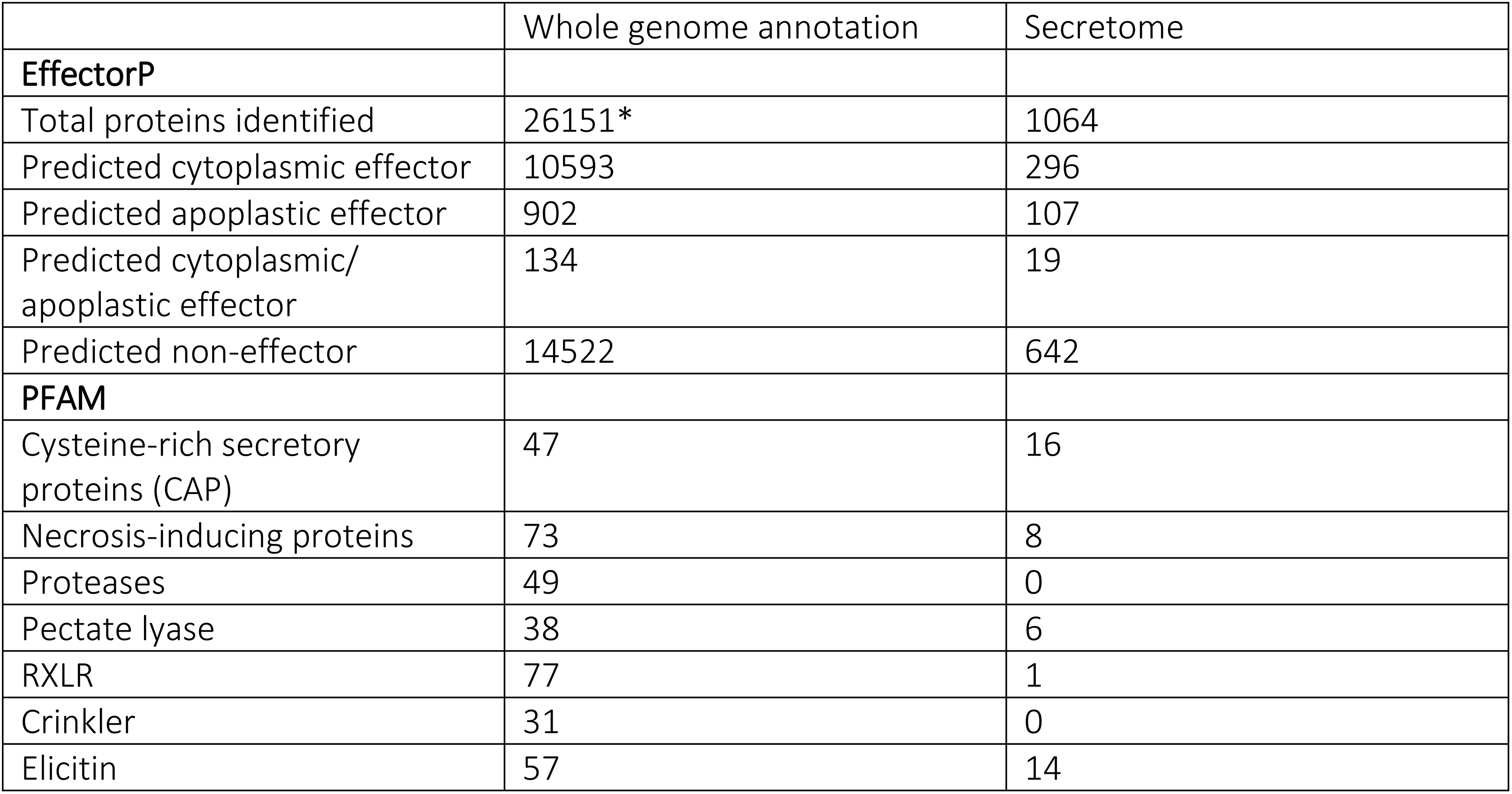
Candidate effector prediction of the *P. cinnamomi* secretome using EffectorP and PFAM.

## Discussion

*Phytophthora cinnamomi* has caused significant losses in crops such as avocado, macadamia and pineapples, in addition to destroying huge areas of natural vegetation and ecosystems world-wide. Not much is known about the molecular mechanisms *P. cinnamomi* infection. Here, we profiled the sub-proteomes of mycelia, zoospores and the secretome using label-free quantitative proteomics to obtain a biochemical snapshot of *P. cinnamomi* that could be used to understand pathogenic success. Although the sub-proteomes were obtained from *in vitro* growth, we present a model of key biochemical processes inferred from protein abundance in zoospore, mycelial and secreted proteins in relation to host tissue (Figure 4).

**Figure 4.**
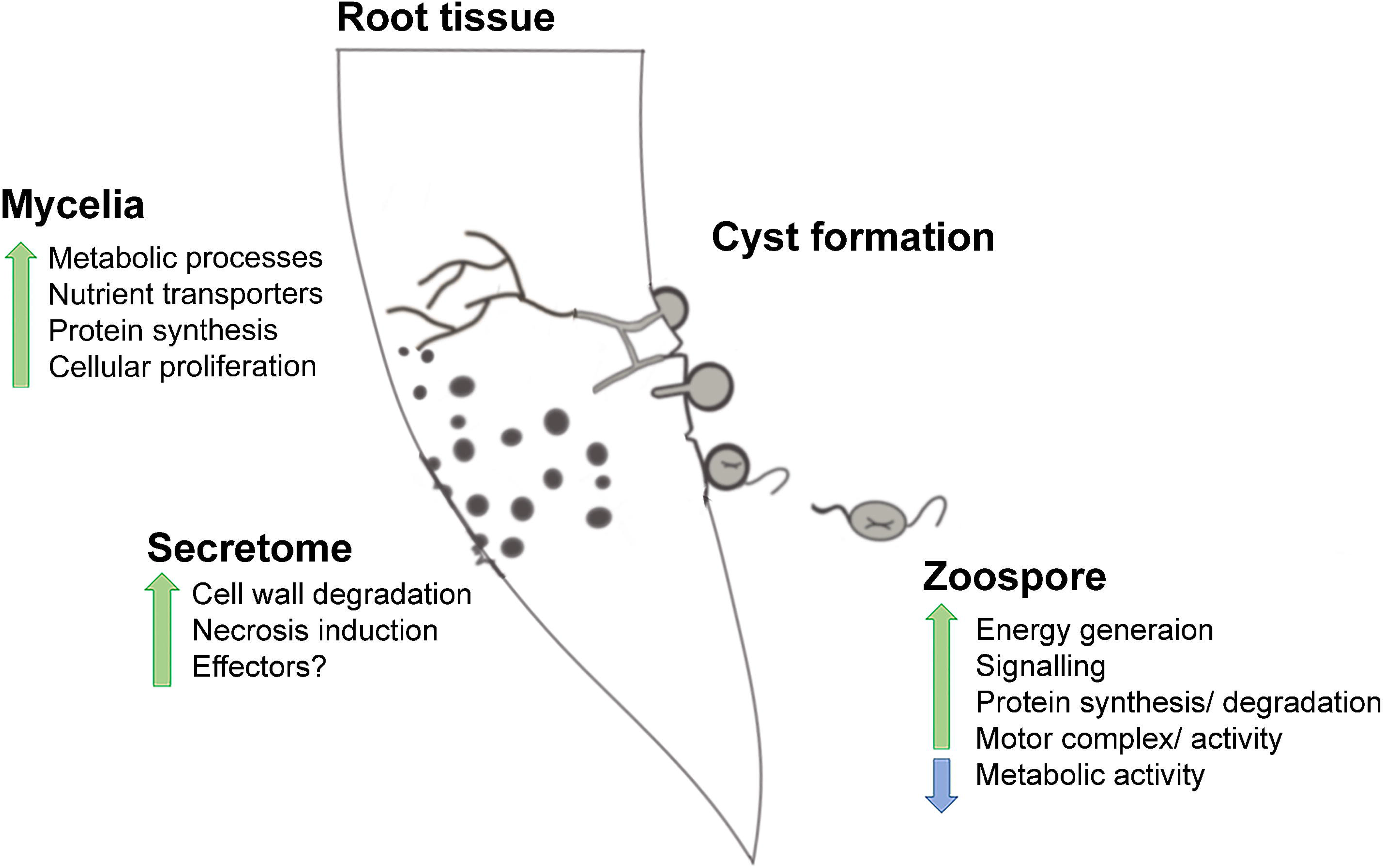
A proposed model of biochemical functions occurring at different sub-proteomes and developmental stages of *P. cinnamomi*.

Mycelia not only grow within root tissue and obtain nutrients through the roots of susceptible plants but are also transmitted to neighbouring plants by direct root-to-root contact ^1^. Several types of transporters are used by oomycetes to acquire nutrients from the extracellular space and host tissue ^18^. 52 transporter genes were identified in the mycelia, showing a priority of nutrient import. This is similar to the mycelial proteomes of *P. pisi*, *P. sojae* and *P. ramorum,* where proteons associated with metabolism, biosynthesis and nutrient transport were enriched or increased in abundance ^19,20^. The enrichment of metabolic processes and cellular reproduction associated proteins in the mycelia indicate that once nutrients have been taken up by the pathogen, they are quickly processed and utilised to facilitate growth (Figure 4).

When conditions favour growth, the mycelia differentiate into sporangia that produce zoospores ^4^. Zoospores have an anterior whiplash and posterior tinsel flagella, which enable them to swim through in moist soil towards potential hosts by tactic gradients ^21,22^. The zoospores are important in the process of infection as they are the first points of contact with the host. The enriched GOs in the zoospores included many associated with energy production such as ATP binding, protein phosphorylation and ATP hydrolysis. Here we hypothesise that there are several needs for zoospores to produce copious amounts of energy. Firstly, zoospore motility requires constant energy production to maintain. Their posterior flagella propels the cells forward and the anterior flagellum is used to steer until they have the opportunity to encyst on the surface of host tissue ^23^. Secondly, once zoospores infect the host, they can switch back to vegetative mycelia where they must derive energy from glycolysis and drive processes that enable the survival of the pathogen within the plant tissue ^20^. The reduced abundance of proteins associated with cellular and lipid metabolism reflects the utilisation of energy generation towards differentiation and motility rather than growth which does not increase until germination of mycelia within host tissue (Figure 4).

The zoospores also showed an increase in RNA binding and modification, protein serine-threonine phosphatase activity, dephosphorylation, protein binding and ubiquitination. This could indicate simultaneous high protein synthesis activity and degradation and may be a response to fast-changing environmental conditions by co-regulating protein synthesis and protein breakdown for proteome adaptation ^24,25^. Enrichment of proteins associated with response to nitrosative stress can be a result of three factors. Nitric oxide (NO) has been shown in pathogenic fungi and oomycetes such as *Phycomyces blakesleeanus* to act as a light sensor molecule during light-mediated sporulation ^15^. Sporulation of *P. cinnamomi* zoospores *in vitro* is light dependent which may use NO as a signalling molecule during this process ^15^. This may also be facilitated by the increase in signalling molecules such as phosphorylation which are abundant in the zoospores. NO is also known to be produced by phytopathogens to aid in early infection such as the oomycete *Bremia lactucae,* where it is important for penetration of the host cell surface ^26^. The nitrosative stress response and nitric oxide dioxygenase activity are important here as even though NO production favours the pathogen during the initial infection, NO accumulation can affect fungal and oomycete growth ^27^. Lastly, an increase in response to nitrosative stress is also utilised in the context of plant production of NO, which is produced by the host as a toxifying defence mechanism ^28^.

Zoospores were also enriched with motor-related proteins such as minus-end-directed microtubule motor activity, cilium assembly, BBSome, dynein complex and kinesin binding proteins. The microtubule is driven by ATP hydrolysis, which was also found to be highly abundant in the zoospores. In oomycetes, dynein complexes are force generated motor proteins that play a crucial role in the movement of the microtubule that drives motility of cilia and flagella ^29^. The BBsome component (analogous to the Baedet-Biedl Syndrome proteins involved in cilia development) in this complex is important in mediating cilia homeostasis in response to stimuli ^30^. The flagella are controlled by motor proteins such as kinesin, which drive the motions of the two types of flagella.

GOs that correspond to proteolysis, cell morphogenesis, tubulin and proteins associated with endocytic recycling were over-represented in the zoospore proteome. When zoospores come into contact with a host, they adhere to the cell surface and form cyst structures ^4^. Proteolytic enzymes can be used in two ways during this process. They are required during cell morphogenesis as the zoospores encyst. Host cell walls are protein-rich therefore proteases are used by the pathogen to degrade cell walls which are subsequently utilised by the mycelia as a nutrient source for growth ^31^. Structural components such as tubulin are also formed as components of the appressorium that penetrates host tissue and allow the pathogen to grow within the plant ^20,32^. This data supports our proposed model that zoospores generate energy to fuel their motility, whilst attempting to dampen the responses of a potential host.

Secreted proteins are of a significant interest due to the localisation mobilisation of effectors into host tissue. Effectors have been identified in many *Phytophthora* species, where they play important roles in infection ^33^. The expression of α- and β-cinnamomin elicitins have been found in various cell types of *P. cinnamomi,* of which β-cinnamomin has been shown to play some role in virulence, however the infection process is a result of a number of effector proteins ^34–37^. The expression of several predicted RXLR effectors has also been demonstrated in *P. cinnamomi* when inoculated onto avocado roots ^38^. Effectors of *P. cinnamomi* have not been fully characterised and effectors at the protein level have not been identified. The secretome profile of *P. cinnamomi* in this study included proteins that contribute to pathogen virulence. This includes enrichment of proteins associated with cell wall modification, carbohydrate metabolism, pectinesterase activity and carbohydrate-binding. These proteins bind and catabolise components of the cell wall that facilitate degradation of host tissue to allow the appressoria to infiltrate and begin infecting the host ^39^. Pectinesterases are proteases that aid in this process by removing esterase from pectin, a major constituent of plant cell walls ^40,41^. The identified pectinesterases catalyse the breakdown of pectin in plant cell walls by binding the active site and lyse the folding structure of the pectin backbone ^42^. The mycelia utilise cell wall degrading enzymes such as glycosidic hydrolases, lectin and acid phosphatase hydrolases as a means of saprophytic in the extracellular space ^43^. Similarly to the secretomes of *P. pisi, P. sojae* and *P. plurivora* several other hydrolases were enriched in the secretome, which can aid host cell disruption and degradation of host defences ^19,40^. In plants, cysteine-type endopeptidases are released in response to pathogen therefore, we hypothesised that hydrolases expressed in *Phytophthora* function to inhibit these proteases from the host ^44–46^

Lupin seedlings were used to test whether the secretome can cause a hypersensitive response *in planta* (Figure 3). Dark lesions on the roots of the seedlings indicate that the secretome contains effector molecules. To gain a deeper understanding of the repertoire of virulence molecules in the secretome, effectorP, PFAM and PHI-Base were used to predict the types of effectors making up the secretome, and look for homology of known effectors (Table 3 and Table 4). 40% of the secretome had predicted cytoplasmic and/or apoplastic effectors. Cysteine-rich proteins were identified in the secretome, which are characteristic of many effectors across oomycete and fungal plant pathogens ^47,48^. Several putative necrosis inducing proteins were also identified along with several elicitins. One of these candidates possess an RXLR domain (o81992|fgenesh1_pm.63_#_55). The RXLR domain is found in some oomycete effector proteins which assists in translocation into host ^49^.

**Table 4.**
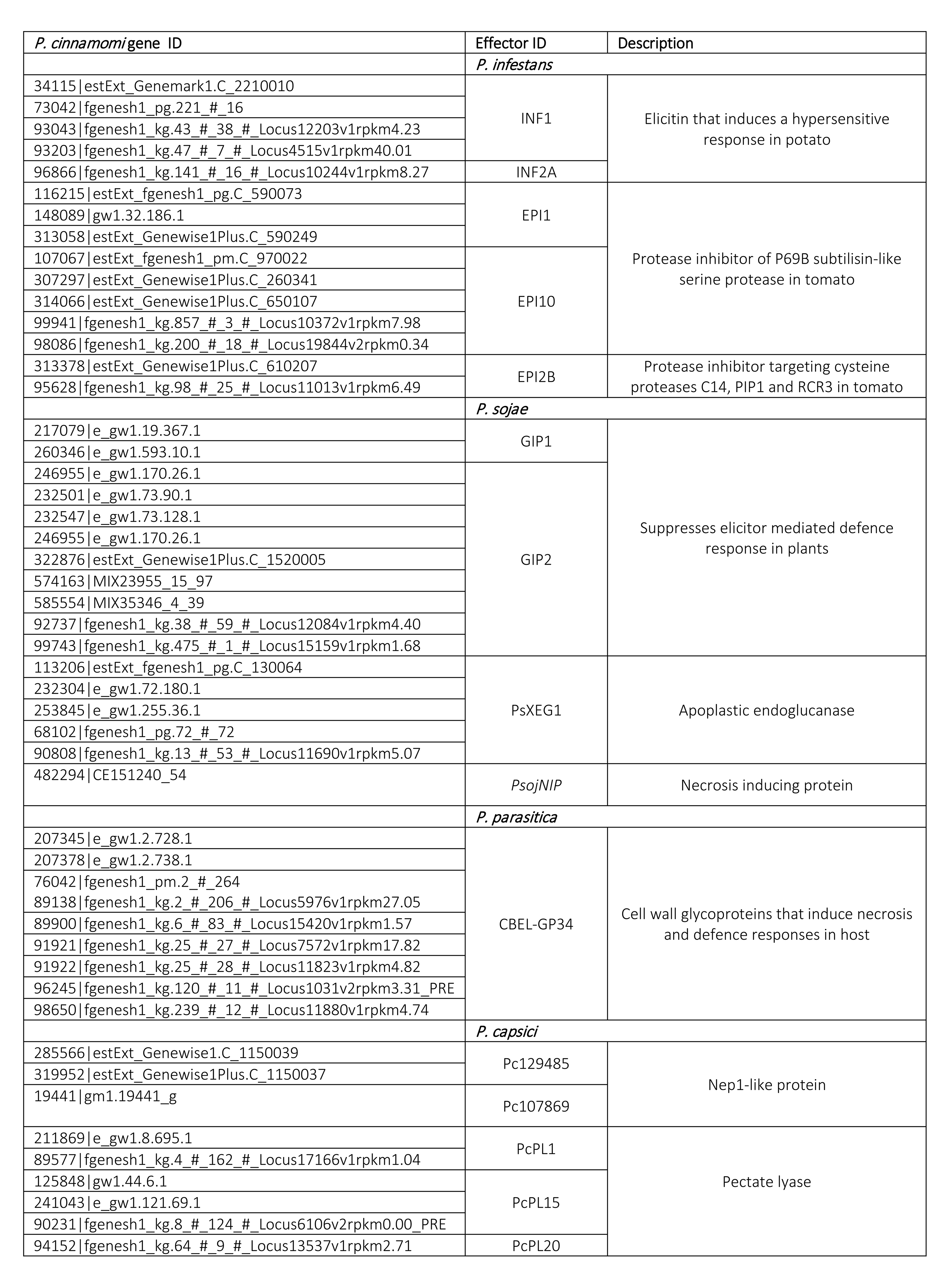
Homology of *P. cinnamomi* secreted proteins to known *Phytophthora* effectors using PHI-Base. A full list of PHI-Base hits to the *P. cinnamomi* secretome is shown in Supplementary material 3.

Searching the set of secreted proteins against the PHI-Base database indicated homology with several *Phytophthora* species (Table 4). Elicitin effectors such as INF1 and INF2A in *P. infestans* have both been shown to cause a hypersensitive response in hosts such as potato ^50,51^. Several protease inhibitors such as EPI1, EPI10 and EPIC2B in *P. infestans,* which block the activity of the proteases, were homologous to proteins detected in the secretome. P69B subtilisin-like serine proteases and EPIC2B are protease inhibitors of cysteine proteases, which are expressed in tomatoes as a defence mechanism against *P. cinnamomi* ^52–54^. Other effectors such as GIP1 and GIP2 in *P. sojae* suppress elicitor mediated defence responses in their hosts ^55^. Homologs to PsXEG1 and CBEL-GP34 were also found, which are adhesive molecules that act on components of the cell walls of hosts in *P. sojae* and *P. parasitica* ^56,57^. Homology of many virulence and effector transcripts between *P. cinnamomi* and *P. infestans* has been demonstrated, including adhesion proteins, hydrolases, proteases and elicitins ^58^. Secretomes identified across many *Phytophthora species* contain a plethora of virulence proteins that contribute in some way to infection of host tissue.

### Concluding remarks

This study provides an in-depth analysis of the protein composition in the mycelia, zoospores and secretome of *P. cinnamomi*. The mycelia were found to be highly metabolically active to support their growth within host tissue and produce molecules that facilitate the breakdown of host tissue that are utilised as nutrients. The biochemical processes in the zoospores are geared towards their motility with an abundance of energy generation to fuel their motility. Here we mapped out the constituents of the motor complex and signalling processes of the zoospores to how they would fit in an infection model. An in-depth dissection of the zoospores has not been previously documented in any *Phytophthora* species. We also provide the first secretome profile of *P. cinnamomi* which includes virulence associated proteins and candidate effectors such as necrosis inducers and elicitins. The discovery of candidate effectors paves the way for future studies and the development of new control measures. This dataset provides a snapshot of the key factors that contribute towards the successful infection of *P. cinnamomi* on its hosts. In future studies, this model can be investigated *in planta* to confirm the biological processes involved in infection and further understand how successful infection is achieved. Due to the emergence of phosphite resistance, there is a need for novel methods of disease management which may be achieved through the discovery of candidate virulence factors such as effectors to aid in the identification of genetic resistance in plants.

## Availability of data and materials

Spectral data used for this study are available at Figshare (DOI: 10.6084/m9.figshare.19161368).

## Acknowledgments

We thank Lars Kamphuis from the Centre for Crop and Disease Management and CSIRO for providing the seed stocks, and Leon Lenzo for his critical feedback on the manuscript. We also thank Owen Duncan for his advice and technical input for the mass spectrometry analysis.

## Abbreviations

MS: Mass spectrometry
LC: Liquid chromatography
GO: Gene ontology
KEGG: Kyoto Encyclopedia of Genes and Genomes
CWDE: cell wall degrading enzymes
BBSome: Baedet-Biedl Syndrome proteins
NO: nitric oxide

